# The Activation of MEK1 by Enhanced Homodimerization Drives Tumorigenesis

**DOI:** 10.1101/229849

**Authors:** Jimin Yuan, Wan Hwa Ng, Zizi Tian, Jiajun Yap, Manuela Baccarini, Zhongzhou Chen, Jiancheng Hu

**Affiliations:** Division of Cellular and Molecular Research, National Cancer Centre Singapore; 11 Hospital Drive, 169610, Singapore; Cancer and Stem Cell Program, Duke-NUS Medical School; 8 College Road, 169857, Singapore; State Key Laboratory of Agrobiotechnology, College of Biological Sciences, China Agricultural University, Beijing 100193, China; Max F. Perutz Laboratories, University of Vienna, Doktor-Bohr-Gasse 9, 1030 Vienna, Austria

**Keywords:** RAF/MEK/ERK kinase cascade, MEK, oncogenic mutations, dimerization, allosteric activity, drug resistance, cancer

## Abstract

Hyperactive RAS/RAF/MEK/ERK signaling has a well-defined role in cancer biology. Aberrant pathway activation occurs mostly upstream of MEK; however, MEK mutations are prevalent in some cancer subsets. Here we show that cancer-related MEK mutants can be classified as those activated by relieving the inhibitory role of helix A, and those with in-frame deletions of β3-αC loop, which exhibit differential resistance to MEK inhibitors in vitro and in vivo. The β3-αC loop deletions activate MEK1 through enhancing homodimerization that can drive intradimer cross-phosphorylation of activation loop. Further, we demonstrate that MEK1 dimerization is required both for its activation by RAF and for its catalytic activity towards ERK. Our study identifies a novel group of MEK mutants, illustrates some key steps in RAF/MEK/ERK activation, and has important implications for the design of therapies targeting hyperactive RAS/RAF/MEK/ERK signaling in cancers.

## Introduction

The RAF/MEK/ERK kinase cascade plays a central role in cell biology, and its hyperactivation results in many human diseases, especially cancers(*1-3*). Genetic alterations that aberrantly activate this kinase cascade in cancers mainly occur in receptor tyrosine kinases (RTKs), RAS, and RAF, while oncogenic mutations of MEK and ERK are rare. However, recent genomic sequencings showed that MEK mutations are highly prevalent in some specific type of cancers such as IGHV4-34^+^ hairy cell leukemia and non-BRAF(V600E) Langerhans cell histiocytosis(*4-7*). To characterize oncogenic MEK mutations and explore how these mutations turn on MEK has important implications for the development of therapeutic inhibitors and for the treatment of MEK-driven cancers.

As a core component of the RAF/MEK/ERK kinase cascade, MEK is a dual specific kinase that transmits a signal from active RAF to phosphorylate ERK(*8, 9*). The regulation of MEK is complex and not yet completely understood. In RAF-driven cancers, active RAF mutants, such as BRAF(V600E), activate MEK by phosphorylating its activation segment, which results in a high level of phospho-MEK(*10*). In contrast, the level of phospho-MEK is much lower in RTK/RAS-driven cancers, although they have similar levels of MEK activity as indicated by comparable phospho-ERK level(*11, 12*). This suggests that wild-type active RAF activates MEK in a different way than BRAF(V600E). In addition, some studies have indicated that AL-autophosphorylation of MEK might also contribute to its activation(*13-15*), adding another layer of complexity to the regulation of MEK activity.

The activity of MEK is regulated not only by the AL-phosphorylation/dephosphorylation but also by its interactions with other components of RAF/MEK/ERK kinase cascade(*16-18*). The heterodimerization of RAF/MEK facilitates MEK activation(*19*), while the MEK/ERK interaction contributes ERK phosphorylation(*20*). Moreover, MEK has been shown to form face-to-face homodimers in which the activation loop of one protomer aligns with the catalytic site of the other(*21*), suggesting a potential intra-dimer transphosphorylation. In addition, MEK also forms heterodimers between its two isoforms, which regulates the duration of ERK signaling(*22*). However, whether and how MEK interactome alterations contribute to hyperactive ERK signaling-driven tumorigenesis remains unkown.

In this study, we have conducted a survey of cancer genomic databases, which leads to the identification of a novel group of cancer-related constitutively active MEK1 mutants with in-frame deletions in the β3-αC loop. These MEK1 mutants exhibit a strong oncogenic potential, but differential sensitivity to MEK inhibitors in clinic therapy or trials. We find that β3-αC loop deletions enhance the homodimerization of MEK1, which drives an intra-dimer transphosphorylation of activation loops between two protomers. Further, we investigated the role of MEK dimerization on the signal-transmission among RAF/MEK/ERK kinase cascade by using these MEK1 mutants with enhanced but differential dimer affinity. Our data show that MEK1 is phosphorylated by active RAF in a dimer-dependent manner and functions as a dimer to phosphorylate ERK1/2, indicating that the dimerization of MEK is critical for its function. Together, this study not only discloses a novel group of oncogenic MEK1 mutants but also uncovers the regulatory mechanism of MEK1 in the RAF/MEK/ERK kinase cascade, and have prominent significance in both drug design and clinic therapy against hyperactive RAS/RAF/MEK/ERK signaling-driven cancers.

## Results

### The MEK mutation spectrum in cancer genomes

To survey and characterize cancer-related MEK mutants, here we interrogated the ICGC (International Cancer Genome Consortium) database, the cBioportal for Cancer Genomics database, and the COSMIC (Catalogue of Somatic Mutations in Cancer) database. Together with those reported in literatures, all MEK mutants in cancer genomes were listed in supplementary table 1, and the MEK mutation spectrum was constructed as in Figure 1A and supplementary Figure 1A. According to our statistic analysis, MEK1 is the dominant isoform of oncogenic alterations, and has four hot spots, which include the negative regulatory helix A (residue 44~61) (I), the αC-β4 loop (II), the β7- β8 loop (III), and the β3-αC loop (IV) (Fig1B-C). It has been shown that the negative regulatory helix A interacts with the kinase domain and stabilizes its inactive conformation, whose disruption will trigger the kinase activity of MEK1 (*23*). According to their structural distributions, we thought that MEK1 alterations/mutations in region I, II, and III activate MEK1 likely through relieving the inhibition of helix A since altered residues are involved in the interaction of helix A with the kinase domain. In contrast, the alterations of β3-αC loop that consist of variable in-frame deletions represent for a novel group of MEK1 mutations, which activate MEK1 via a distinct mechanism.

**Figure 1.**
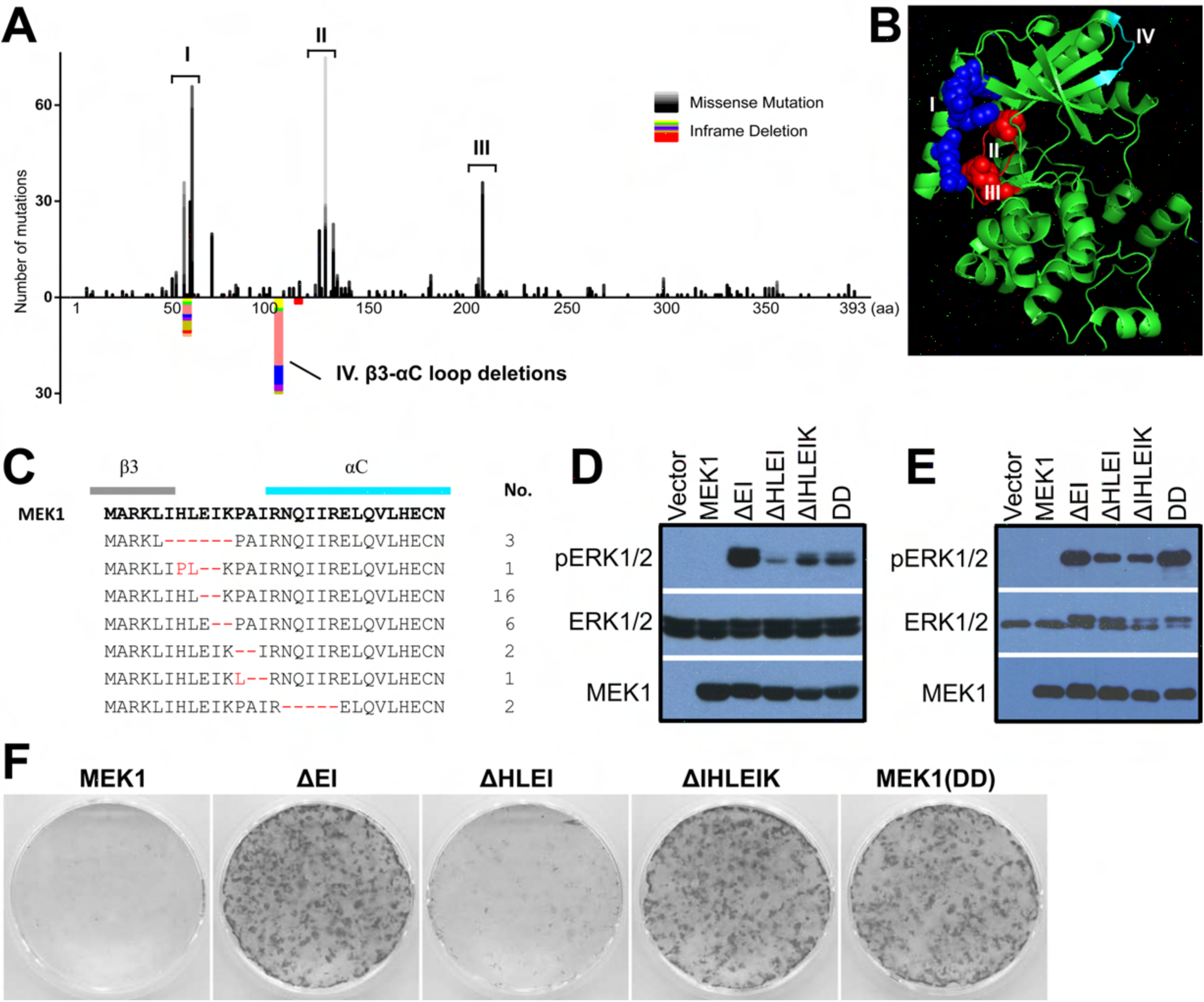
The β3-αC loop deletion defines a novel group of MEK mutations in cancer genomes. (A) Cancer-related MEK1 mutation spectrum. Cancer-related MEK1 mutations have four hotspots: (I) the inhibitory helix A, (II) the αC-β4 loop, (III) the β7- β8 loop, and (IV) the β3-αC loop. (B) Schematic diagram showing the mutation hotspots of MEK1. The frequently mutated-residues in helix A are colored with blue, and those in hotspot II and III are colored with red. All of these residues in hotspot I, II, and III involve in the interaction of helix A with the kinase domain. The hotspot IV is colored with cyan. (C) Alignment of MEK1 mutants with β3-αC loop deletions. (D-E) MEK1 mutants with β3-αC loop deletions are constitutively active. D, MEK1 mutants activate ERK signaling when expressed in 293T cells. E, MEK1 mutants phosphorylate ERK in vitro upon purification from 293T transfectants. (F) MEK1 mutants with β3-αC loop deletions transform immortalized MEFs. The stable fibroblast cell lines expressing MEK1 mutants were constructed and the foci formation assay was carried out as described before(*27, 29*).

### MEK1 mutants with in-frame deletions of β3-αC loop are oncogenic and have differential resistance to MEK inhibitors

To characterize MEK1 mutants with in-frame deletions of β3-αC loop, we measured their activity in vivo and in vitro. As shown in Figure 1D-E and supplementary Figure 1B-D, these mutants activate ERK1/2 when expressed in 293T cells or immortalized fibroblasts with or without RAFs, or upon purification from 293T transfectants, suggesting that they have constitutive activity independent of upstream stimuli. Further, all MEK1 mutants except MEK1(ΔHLEI) that has the lowest activity exhibit a strong oncogenic potential, which transform cells independently of RAFs when expressed in immortalized fibroblasts (Fig 1F and S1E-F) and induce fibroblastomas when these transformed fibroblasts were xenografted into NOD-SCID mice (data shown below). Together, these data indicated that MEK1 mutants with in-frame deletions of β3-αC loop could function as cancer drivers.

We next determined whether MEK inhibitors applied in therapy or in clinic trials(*24*) could be used to target oncogenic MEK1 mutants with variable deletions of β3-αC loop. By using 293T transfectants and stable fibroblast lines expressing these MEK1 mutants, we found that, among all inhibitors that tested in this study, Tramentinib was the most effective in blocking the activity of MEK1(ΔEI), while MEK1(ΔIHLEIK) was resistant to all inhibitors (Fig 2A and S2A-C). To further evaluate the ability of Tramentinib to target MEK1 mutants with β3-αC loop deletions in vivo, we here constructed xenografted melanomas by using MeWo melanoma cell lines that stably express these MEK1 mutants. Together with xenografted fibroblastomas above, we found that all tumors harboring MEK1(ΔIHLEIK) had a robust resistance to Tramentinib treatment in contrast to those harboring MEK1(ΔEI) or MEK1(DD) (Fig 2B-D and S2D-G).

**Figure 2.**
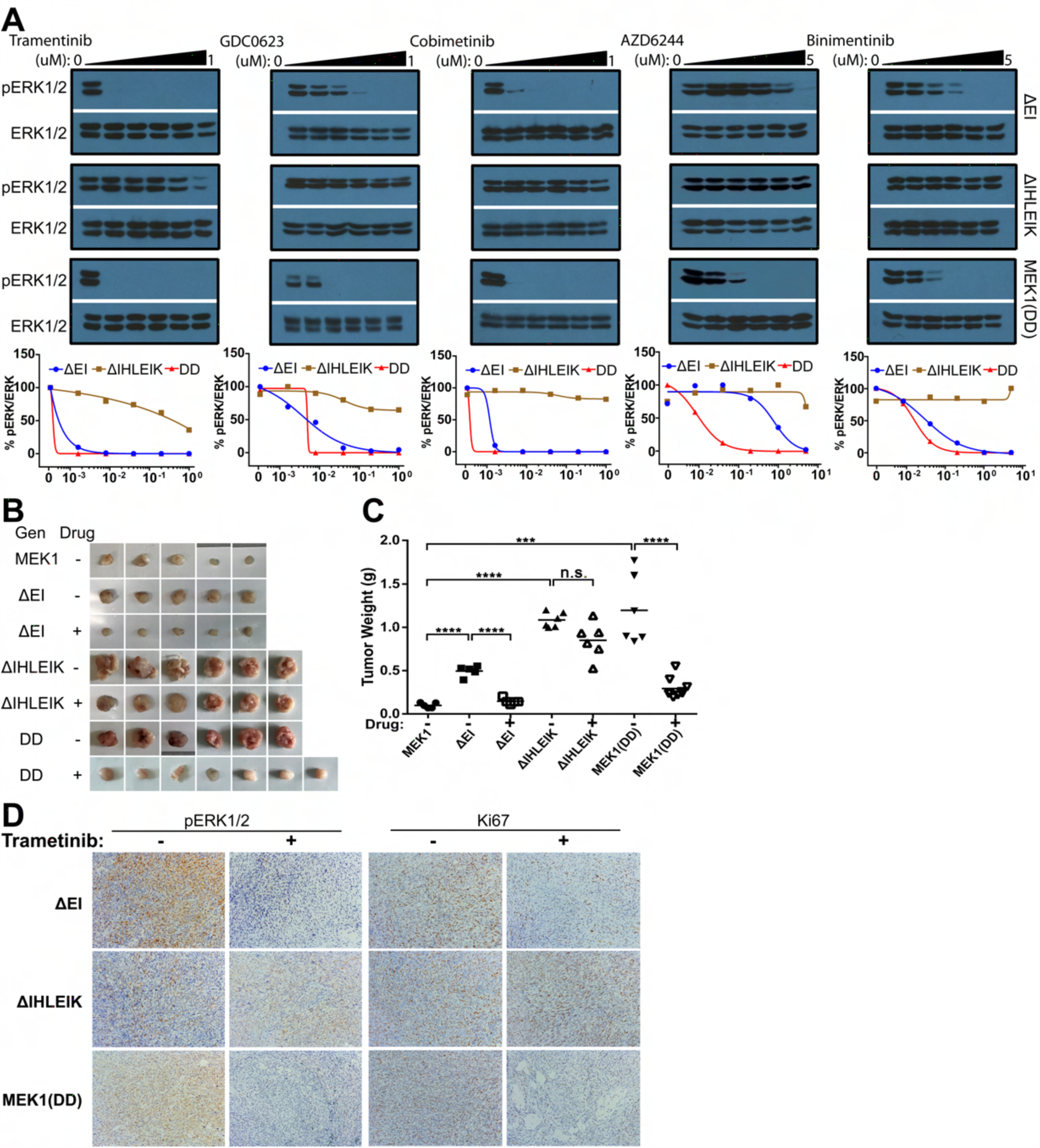
MEK1 mutants with β3-αC loop deletions exhibit a differential inhibitor resistance in vitro and in vivo. (A) MEK1 mutants with β3-αC loop deletions have different sensitivity to MEK inhibitors when expressed in fibroblasts. Fibroblasts stably expressing different MEK1 mutants were treated with MEK inhibitors as indicated, and phospho-ERK1/2 was probed by immunoblot and quantified by using Image J. (B-D) Fibroblastomas induced by MEK1 mutants with β3-αC loop deletions have different response to MEK inhibitor treatment. B-C, Fibroblasts expressing different MEK1 mutants were subcutaneously injected NOD-SCID mice which will be treated with or without tramentinib as described in Materials and Methods, and xenograft tumors were harvested and weight. D, Immunohistochemistry staining analysis of xenograft tumors from B. ERK1/2 activity and cell proliferation were indicated respectively as phosphor-ERK1/2 and Ki67 stainings.

### In-frame deletions of β3-αC loop activate MEK1 through enhancing homodimerization

In-frame deletions of β3-αC loop define a new group of oncogenic MEK1 mutations. However, how these alterations trigger the kinase activity of MEK1 remains unknown. Recently, we and other groups found that variable β3-αC loop deletions could activate RAF kinases through promoting homodimerization(*25,26*) (Yuan and Hu, unpublished data). Here we assumed that these MEK1 mutants are activated via a similar manner though it has been shown that, different from the side-to-side dimer of RAF kinases, MEK1 forms a face-to-face dimer(*21*). To testify this notion, we firstly measured the dimer affinity of MEK1 mutants with variable β3-αC loop deletions by using PAGE with low SDS(0.1%) and microscale thermophoresis (MST) methods. As shown in Figure 3A and S3A, MEK1 mutants with in-frame β3-αC loop deletions have elevated dimer stability/affinity with MEK1(ΔIHLEIK) > MEK1(ΔEI) > MEK1(DD) > MEK1(ΔHLEI) > WT. A compound mutation that includes N78G, V224G, F311A, and L314A (called as GGAA below) in the dimer interface(*21, 22*), dramatically decreased both the dimer stability/affinity and the activity of MEK1 mutants with β3-αC loop deletions (Fig 3B and S3B), which further supports our hypothesis.

**Figure 3.**
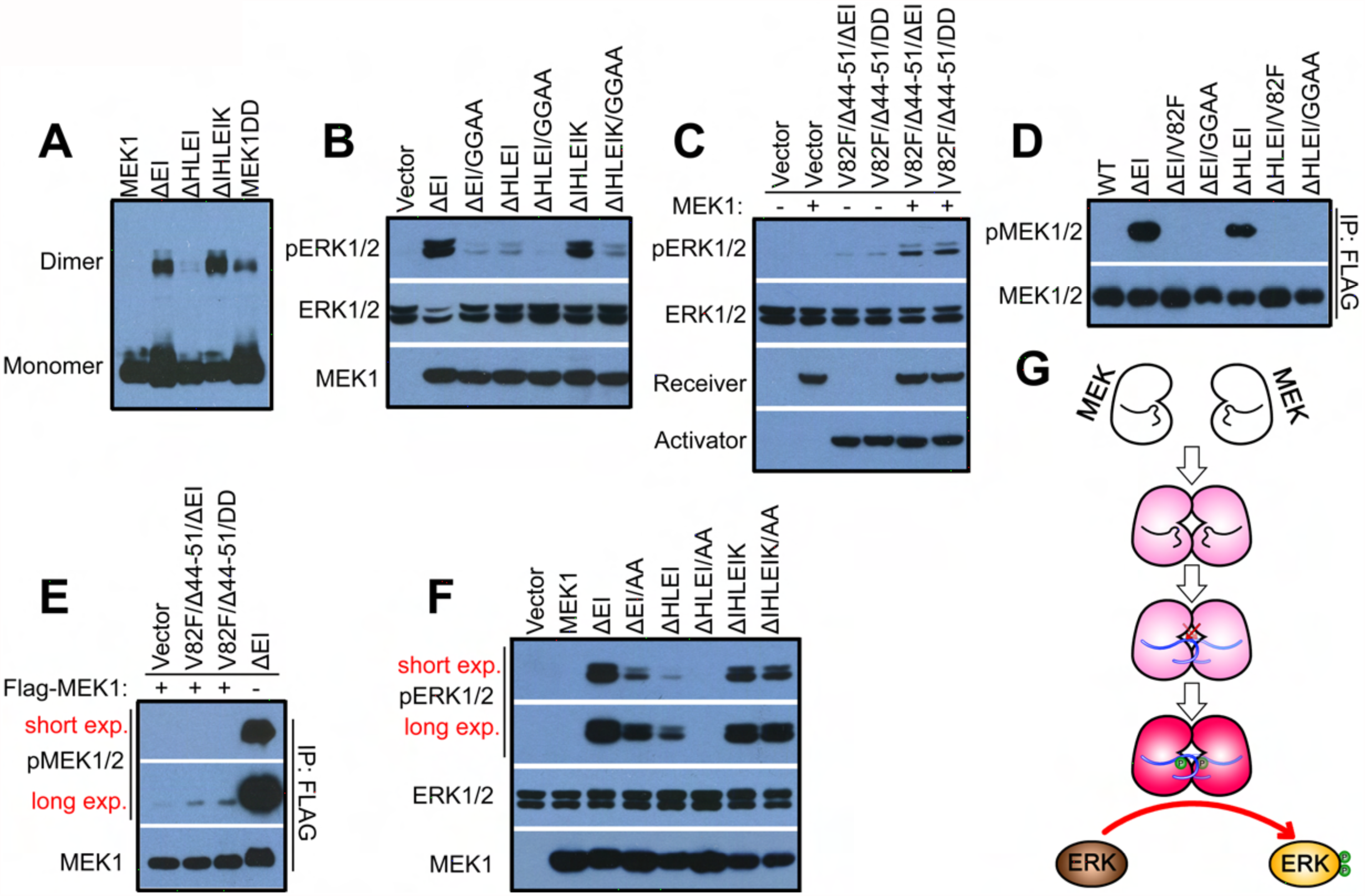
Molecular mechanism that underlies the activation of MEK1 by variable β3-αC loop deletions. (A) The β3-αC loop deletions promote the homodimerization of MEK1. MEK1 mutants were expressed in 293T cells and their oligomerization were detected by PAGE with low SDS (0.1%). (B) The compound mutation (GGAA) that disrupts the dimer interface blocks the activity of MEK1 mutants with β3-αC loop deletions. MEK1 mutants were expressed in 293T cells, and their activity was measured by phospho-ERK1/2 immunoblot. (C) Kinase-dead MEK1 is able to function as allosteric activator to stimulate wild-type MEK1. C-spine fused MEK1 mutants were expressed respectively with WT MEK1 in 293T cells, phospho-ERK1/2 was measured by immunoblot. (D-F) The activation of MEK driven by homodimerization requires an intra-dimer cross-phosphorylation of AL between two protomers. D, MEK1(ΔEI) and MEK1(ΔHLEI) auto-phosphorylate their AL, which is blocked C-spine fusion. MEK1 mutants were purified from 293T transfectants and the phospho-AL was detected by immunoblot. E, the AL of MEK1 receiver is not phosphorylated when coexpressed with allosteric activators. The wild-type MEK1 was co-expressed with kinase-dead allosteric MEK1 mutants, and its AL-phosphorylation was detected by immunoblot upon immunoprecipitation. F, The non-phosphorylatable AL-mutation decreases the activity of MEK1(ΔEI) and MEK1(ΔHLEI). MEK1 mutants were expressed in 293T cells, and phospho-ERK1/2 in transfectants was detected by immunoblot. (G) A model of homodimerization-driven MEK activation. Homodimerization helps MEK assembling an active conformation, which in turn induces a cross-phosphorylation of AL between two protomers. The AL-phosphorylation fully activates MEK dimer and stabilizes its active conformation. Then MEK dimer with phospho-AL catalyzes the phosphorylation of ERK AL.

The activity of MEK1 is triggered by the enhanced homodimerization, suggesting that it has both allosteric and catalytic functions as RAF kinases do. To examine the allosteric activity of MEK1 by protein kinase co-activation assay(*27-33*), we created kinase-dead allosteric activators by deleting the negative regulatory helix A (ΔE44-E51) and fusing the catalytic spine (V82F mutation) in MEK1 mutants with variable deletions of β3-αC loop (Fig S3C). MEK1(ΔEI/helix A^-^/V82F) was the only mutant capable of activating wild-type MEK1 when coexpressed in 293T cells, and its allosteric activity was blocked by the GGAA mutation in dimer interface (Fig S3D-E). The subtle allosteric activity of MEK1(ΔHLEI/ helix A^-^/V82F) and MEK1(ΔIHLEIK/ helix A^-^/V82F) might arise from their weak ability to dimerize with receiver (Figure S3F). Further, we examined whether a compound MEK1 mutant mimicking active MEK1 with AL-phosphorylation has allosteric activity. As shown in Figure 3C, the MEK1 mutant containing the helix A^-^, the V82F, and the DD (S218D, S221D), could stimulate the activity of wild-type MEK1 as well as MEK1(ΔEI/ helix A^-^/V82F) when co-expressed in 293T cells. Together, these data demonstrate that wild-type MEK1 has an ability of allosteric activator once activated.

The AL-phosphorylation is an indicator of MEK fully activation(*8*). To explore the molecular basis of MEK1 activation by enhanced homodimerization, we checked the AL-phosphorylation status of MEK1 mutants with variable deletions of β3-αC loop, and found that the AL was phosphorylated in MEK1(ΔEI) and MEK1(ΔHLEI) but not in MEK1(ΔIHLEIK). Moreover, their AL-phosphorylation was blocked by either catalytic spine fusion or dimer interface disruption (Fig 3D and S3G). This data suggested that MEK1 was turned on by homodimerization and then auto-phosphorylated its AL, and that MEK1(ΔIHLEIK) might mimic an intermediate locked in a transitional status. To further determine whether the AL-autophosphorylation occurs in cis or trans, we purified the receiver from MEK1 coactivation transfectants and found that its AL was barely phosphorylated (Fig 3E), suggesting that the homodimerization drives a intradimer cross-phosphorylation of two protomers. This finding was also supported by that the non-phosphorylatable AL-mutation (S217A/S221A) reduces the activity of MEK1(ΔEI) and MEK1(ΔHLEI) by 60~70% but not that of MEK1(ΔIHLEIK) (Fig 3F). Overall, our data demonstrated that in-frame deletions of β3-αC loop activate MEK1 via homodimerization-driven cross-phosphorylation (Fig 3G).

### MEK1 transduces a signal from active RAF to activate ERK in a dimer-dependent manner

The interactions among different components play a crucial role in the signal transmission of RAF/MEK/ERK kinase cascade(*16-18*). Recently, we demonstrated that active RAF kinases function as a dimer to phosphorylate MEK and the MEK-binding ability of both RAF protomers is required for this process (Yuan and Hu, unpublished). To further understand the molecular basis of this process, here we determined whether MEK is phosphorylated by active RAF as a dimer. Thus, we coexpressed BRAF(V600E) or K-RAS(G12V) with wild-type MEK1, or mutants with impaired/enhanced dimer, and found that both BRAF(V600E) and K-RAS(G12V) induced the AL-phosphorylation of wildtype MEK1 and dimeric MEK1(ΔIHLEIK) but not of dimer-impaired MEK1(GGAA) mutant, though the level of AL-phosphorylation differed (Fig 4A). This suggests that MEK dimerization is indispensable for its AL-phosphorylation by active RAF. Since the activation of MEK requires its heterodimerization with RAF, and the heterodimer interface on MEK mostly overlaps with its homodimer interface(*19*), it is possible that the impaired AL-phosphorylation of MEK1(GGAA) induced by BRAF(V600E) or K-RAS(G12V) results from a loss of heterodimerization with RAF. To investigate this possibility, we examined the interaction of MEK1 mutants with BRAF(V600E) by co-immunoprecipitation, and found that both MEK1(GGAA) and MEK1(ΔIHLEIK) showed a reduced complex formation with BRAF(V600E) compared to wild-type MEK1 (Fig 4B), indicating that the impaired AL-phosphorylation of MEK1(GGAA) is not related to loss of heterodimerization with RAF. In addition, the waeker AL-phosphorylation of MEK1(ΔIHLEIK) induced by K-RAS(G12V) but not by BRAF(V600E) compared to wild-type MEK1 suggested that the heterodimerization of MEK1 with RAF is important for its activation by wild-type RAF. Together, these data indicate that the dimerization of MEK1 is required for its phosphorylation by active RAF.

**Figure 4.**
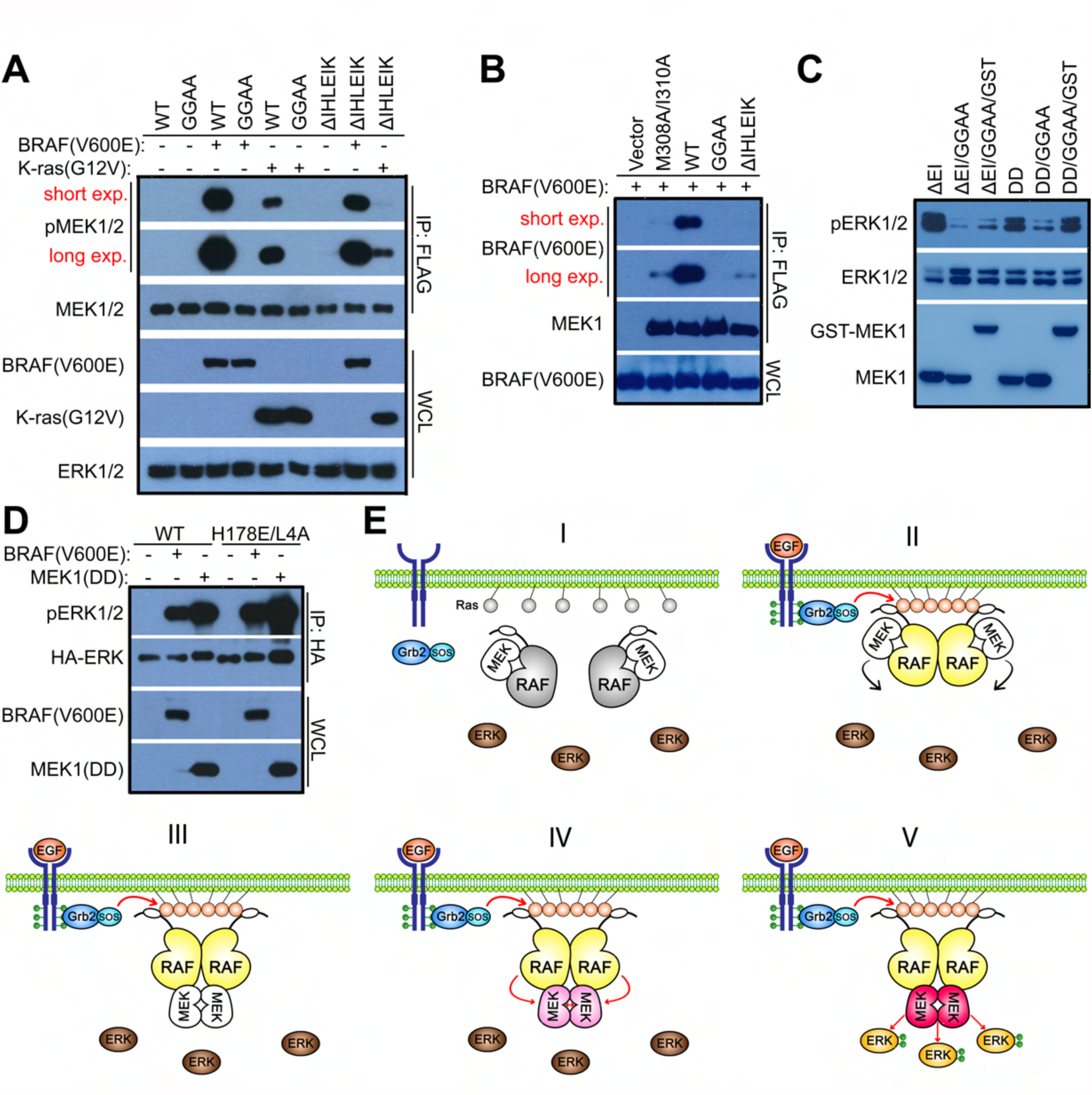
The dimerization of MEK is critical for its phosphorylation by active RAF and for its catalytic activity towards ERK. (A-B) The phosphorylation of MEK by active RAF requires its dimerization. A, Wild-type MEK1 and the dimer-enhanced MEK1 mutant but not the dimer-impaired MEK1 mutant are phosphorylated by active RAF. MEK1 mutants were co-expressed with either BRAF(V600E) or K-ras(G12V) in 293T cells, and their AL-phosphorylation is detected by immunoblot upon immunoprecipitation. B, Both dimer-enhanced and dimer-impaired MEK mutants barely bind to active RAF. MEK1 mutants were co-expressed respectively with BRAF(V600E) and their dimerization was detected by co-immunoprecipitation. MEK1(M308/310A) that has an impaired ability to dimerize with RAF kinase serviced as a control. (C) Active MEK functions as a dimer to activate ERK. The disruption of dimer interface inhibits the activity of MEK1(DD), which can be rescued by GST fusion. In contrast, this does not occur in MEK1(AEI), since it is triggered by enhanced-dimerization. MEK1 mutants were expressed in 293T cells, and their activity was measured by phospho-ERK1/2 immunoblot. (D) Both wild-type and dimer-impaired ERK can be phosphorylated by MEK. ERK2 mutants were expressed with either MEK1(DD) or BRAF(V600E) in 293T cells, purified by immunoprecipitation, and their AL-phosphorylation was detected by immunoblot. (E) A model for the activation of RAF/MEK/ERK kinase cascade. I. In quiescent cells, RAF/MEK forms a face-to-face heterodimer in cytosol. II. Upon stimulation, RAF/MEK heterodimers are recruited to the plasma membrane by active RAS through the RBD domain of RAF, where they form a transient tetramer through the side-to-side RAF dimerization. III. The side-to-side RAF dimerization not only turns on RAF but also loosens its face-to-face heterodimerization with MEK, and therefore facilitates the homodimerization of MEK on the surface of RAF dimer. IV. Active RAF dimer activates the MEK dimer docking on its surface by phosphorylating its AL, and MEK dimer can also activate itself through homodimerization-driven AL-auto-phosphorylation. V. Active MEK dimer docking on or releasing from the surface of RAF dimer phosphorylates ERK.

We next investigated whether active MEK functions as a dimer and requires dimeric ERK as substrate for phosphorylation as active RAF does. Indeed, the disruption of dimer interface by GGAA mutation impairs the activity of MEK1(DD) toward ERK which can be restored by GST fusion (Fig 4C), suggesting that active MEK really acts as a dimer. In contrast, wild-type and monomeric ERK were phosphorylated equally when co-expressed with BRAF(V600E) or MEK1(DD) in 293T cells (Fig 4D), indicating that the dimerization of ERK is dispensable for its activation by MEK.

## Discussion

In this study, we systematically characterized cancer-related MEK mutations, and found that MEK mutants could be classified as two groups: (1) mutants in which the inhibitory interaction between helix A and kinase domain is relieved, and have been shown sensitive to MEK inhibitors(*5*, *6*, *34*, *35*); (2) mutants with in-frame deletions of β3-αC loop, which are turned on through enhanced homodimerization and have differential resistance to MEK inhibitors. This finding provides a guideline for the targeted therapy of MEK-driven cancers.

MEK1 mutants with variable deletions of β3-αC loop exhibit enhanced but variable dimer affinity, different degrees of AL-phosphorylation, and differential drug resistance. Previous studies have shown that, although the activity of ERK in both BRAF(V600E)- and Ras mutant-driven cancers is comparable, the AL-phosphorylation of MEK and the sensitivity to MEK inhibitors are quite different(*10-12*), raising the possibility that the inhibitor sensitivity correlates with the AL-phosphorylation status of MEK. Our study excludes this possibility in MEK1 mutants with variable deletions of β3-αC loop (data not shown), but whether their different inhibitor sensitivity results from their differential dimer affinity requires further study.

The activation of the RAF/MEK/ERK kinase cascade is a complex process involving physical interactions among its components(*16-18*). This process is initiated by RAF dimerization, which not only turns on RAF but also facilitates the activation of MEK. Then MEK could be activated by RAF-mediated AL-phosphorylation, but also by homodimerization-driven AL-auto-phosphorylation. Like RAF kinases, the dimerization of MEK is required for both its activation and catalytic function. Together with previous findings, we propose the following model for the activation of the RAF/MEK/ERK kinase cascade (Fig 4E): (1) In quiescent cells, RAFs form face-to-face heterodimers with MEKs. (2) Upon stimulation, active RAS (also dimers) recruit RAF/MEK heterodimers to the plasma membrane through the RBD domain of RAFs. There, RAF/MEK heterodimers form transient tetramers by the back-to-back dimerization of RAFs. (3) The back-to-back dimerization of RAFs activates RAFs by inducing cis-autophosphorylation of the AL, and on the other hand, loosens RAF/MEK heterodimers and facilitates the assembly of MEK homodimers on their surface. (4) Active RAF dimers phosphorylate one or both protomers of MEK dimers docking on their surface, and the phospho-MEK protomers can also cross-phosphorylate the other protomers in the context of MEK homodimers. (5) Once both protomers are phosphorylated, MEK dimers can be released from the RAF dimers and phosphorylate ERKs. Semi/non-phosphorylated active MEK dimers docking on the surface of active RAF dimers can also directly phosphorylate ERKs. Constitutively active RAF mutants including BRAF(V600E) also activate MEK in a dimer-to-dimer manner. According to this model, allosteric inhibitors that disrupt RAF-MEK and MEK-MEK dimerization would have a high efficacy to block hyperactive ERK signaling in cancers harboring genetic alterations in MEK or upstream of MEK, and be ideal next-generation drugs for cancer therapy.

## Materials and Methods

### Antibodies, Biochemicals, Cell lines, and Plasmids

Antibodies used in this study include: anti-phosphoERK1/2, anti-phosphoMEK1/2, and anti-MEK1/2 (Cell Signaling Technology); anti-FLAG (Sigma); anti-HA (Novus Biologicals); anti-ERK1/2 (AB clonal); anti-Ki67 (Abcam); and HRP-labeled secondary antibodies (Jackson Laboratories). Trametinib, GDC0623, Cobimetinib, AZD6244, and Binimentinib were purchased from Medchemexpress. All other chemicals were obtained from Sigma. Wild-type, BRAF^-/-^ and CRAF^-/-^ fibroblasts were generated in previous study(*36*). MeWo melanoma cell line was obtained from ATCC. Plasmids encoding RAS, RAF, MEK, ERK, and their mutants were constructed by Gibson assembly. pCDNA3.1(+) vector (Invitrogen) was used for transient expression; retro- or lenti-viral vectors (Clontech) for stable expression; and pET-28a (Novagen) for bacterial expression.

### Protein expression and Purification

6xhis-tagged ERK2(K52A) was expressed in BL21(DE3) strains and purified by using Nickel column (Qiagen) as described before(*37*). FLAG-tagged MEK1, MEK1(ΔEI), MEK1(ΔIHLEIK) were expressed in 293T cells and purified by using anti-FLAG affinity gel and 3XFLAG peptide (Sigma) and following manufacturer’s protocol.

### Cell Culture, Transfection, and Transduction

All cell lines were maintained in DMEM medium with 10% FBS (Hyclone). Cell transfection were carried out by using the Biotool transfection reagent. To generate stable cell lines that express MEK1 mutants, viruses were prepared and applied to infect target cells according to our previous studies(*37, 38*). Infected cells were selected by using antibiotics.

### Immunoprecipitation, *In Vitro* Kinase Assay, and Western Blotting

Immunoprecipitations were performed as described previously(*27-29*). Briefly, whole-cell lysates were mixed with either anti-HA, or anti-FLAG beads (Sigma), rotated in cold room for 60 min, and washed three times with RIPA buffer. For *in vitro* kinase assays, the immunoprecipitants were washed once with kinase reaction buffer (25 mM HEPES, 10 mM MgCl2, 0.5 mM Na3VO4, 0.5 mM DTT, pH 7.4), then incubated with 20μl kinase reaction mixture (2 ug substrate and 100 mM ATP in 20μl kinase reaction buffer) per sample at room temperature for 10 min. Kinase reaction was stopped by adding 5μl per sample 5XLaemmli sample buffer. Immunoblotting was carried out as described before(*37*).

### Foci formation assay

Immortalized MEFs infected with retroviruses encoding target proteins were plated at 5×10^3^ cells per 60mm dish, and fed every other day. 12 days later, cells were fixed with 2% formaldehyde and stained with Giemsa solution (Sigma).

### Animal studies

For xenograft experiments, female NOD/SCID mice (6~8 weeks) were subcutaneously injected with 3x10^6^ cells per mice in 1:1 matrigel (Corning). Tumor volumes were monitored by digital calipers twice a week and calculated using the formula: volume= (width)^2^ × length/2. Tramentinib was administered orally (2mg/kg) every other day when tumors reached an average volume of ~50-60 mm^3^. At the experiment endpoint, mice were euthanized and tumors were harvested for *ex vivo* analysis.

### Immunohistochemistry staining

Tumors were fixed in 10% buffered formalin overnight and embedded according to standard procedures. Tumor sections were cut to 4um thickness, mounted on glass slides, and air-dried at room temperature. After antigen retrieval, tumor sections were stained with antibodies and then with hematoxylin. Images of tumor sections were taken with a bright light microscope at X10.

## Author Contributions

J.Y. and J.H. designed the study; J.Y. and J.H. searched databases/literatures for MEK mutations in cancer genomes; M.B. prepared RAF-knockout cell lines; J.Y., W.H.N, Z.T., J.J.Y., and J.H. carried out molecular biology, biochemistry, and cell biology experiments; J.Y. constructed mouse xenograft models and performed immunohistology analysis; M.B., Z.C. and J.H. supervised all experiments and interpreted experimental data; J.H. wrote the manuscript; M.B. revised manuscript; and all authors commented and approved the manuscript.

## Acknowledgement

We thank Dr. Hui Kam Man, Dr. Kanaga Sabapathy, Dr. Paula Lam, Dr. David Virshup, Dr. Wang Mei, and their laboratories for their help in experimental approaches and comments on this manuscript. We also thank Dr. Andrey Shaw and Dr. Susan Taylor for their assistances. This study is supported by NCCRF startup grant (NCCRF-SUG-JH), NCCRF bridging grant (NCCRF-YR2016-JUL-BG1), NMRC seeding grants (NCCSPG-YR2015-JUL-14 and NCCSPG-YR2016-JAN-17), Duke-NUS Khoo Bridge Funding Award (Duke-NUS-KBrFA/2017/0003), Asia Fund Cancer Research (AFCR2017/2019-JH) and SHF Research Grant (SHF/FG692S/2016). The authors declare no potential conflicts of interest.

